# Fatigue >12 weeks after coronavirus disease (COVID) is associated with reduced reward sensitivity during effort-based decision making

**DOI:** 10.1101/2024.11.04.621806

**Authors:** Judith M. Scholing, Britt I.H.M. Lambregts, Ruben van den Bosch, Esther Aarts, Marieke E. van der Schaaf

**Affiliations:** Donders Institute for Brain, Cognition and Behaviour, Radboud University, Nijmegen, The Netherlands; Tilburg University, Department of Cognitive Neuropsychology, Tilburg, The Netherlands

**Author notes:** ^#^These authors contributed equally (Esther Aarts, Marieke E. van der Schaaf). **Corresponding author**: Judith M. Scholing, **Address:** Donders Institute for Brain, Cognition and Behaviour, Kapittelweg 29, 6525 EN Nijmegen.

**Keywords:** COVID-19, post-COVID, chronic fatigue, motivation, effort-based decision making

## Abstract

**Background:** Fatigue and depressive mood is inherent to acute disease, but a substantial group of people report persisting disabling fatigue and depressive symptoms long after a COVID infection. Acute infections have shown to change decisions to engage in effortful and rewarding activities, but it is currently unclear whether fatigue and depressive symptoms similarly affect decision making during acute and persistent phases of a COVID infection. Here, we investigated whether fatigue and depressive mood are associated with altered weighting of effort and reward in decision-making during different timepoints after COVID infection.

**Methods:** We conducted an online cross-sectional study between March 2021 and March 2022. 242 Participants (18-65 years) with COVID <4 weeks ago (n=62), COVID >12 weeks ago (n=81), or no prior COVID (self-reported) (n=90) performed an effort-based decision-making task, in which they decided whether they wanted to exert physical effort (ticking boxes on screen, 5 levels) for reward (money to be gained in a voucher-lottery, 5 levels). State fatigue and depressive mood was measured by the Profile of Mood States (POMS) prior to the task. We used multilevel binomial regression analysis to test whether fatigue and depressive mood were related to acceptance rates for effort and reward levels and whether this differed between the groups.

**Results:** Compared with no COVID and COVID <4 weeks groups, the COVID >12 weeks group reported higher state fatigue scores (mean±SD: 20±7 vs. 14±7 and 12±6 POMS-score, respectively; both p<0.001) and was less sensitive to rewards (Reward*Group: OR: 0.35 (95%CI 0.20, 0.62), p<0.001 and OR: 0.38 (95%CI 0.20, 0.72), p=0.003). In the COVID >12 weeks group, fatigue was more negatively associated with reward sensitivity compared with the COVID <4 weeks group (Reward*Fatigue*Group: OR 0.47 (95%CI 0.25, 1.13), p=0.022) and the no COVID group (Reward*Fatigue*Group: OR 0.48 (95%CI 4.01, 0.92), p=0.029). No group differences were observed for the relationship between fatigue and effort sensitivity. No group differences were observed for the relationship between depressive mood and effort or reward sensitivity. Higher age, lower BMI, unhealthy lifestyle, and worrying during the acute phase of COVID each predicted decreased reward sensitivity in the >12 weeks group (Age*Reward: OR 0.30 (95%CI 0.19, 0.48), p<0.001; BMI*Reward: OR 1.43 (95%CI 1.01, 2.00), p=0.047); Lifestyle*Reward: OR 1.50 (95%CI 1.06, 2.14), p=0.022; Worrying*Reward: OR 0.59 (95%CI 0.38, 0.94), p=0.025, respectively).

**Conclusion:** The finding that fatigue is related to lower reward sensitivity >12 weeks after COVID, suggesting potential reward deficits in post-covid fatigue. These findings are in line with previous observations that long-term inflammation can induce dysregulations in neural reward processing, which should be further investigated in future studies.

**Highlights:**

- We tested if fatigue and mood were related to altered decision making post-COVID
- Participants post-COVID >12wks ago were more fatigued and less reward sensitive
- Post-COVID-related fatigue was associated with reduced reward sensitivity
- Post-COVID-related depressive mood was not associated with altered decision making
- Higher age, unhealthy lifestyle, and worrying predicted reward deficits

## Introduction

Flu-like symptoms such as fever, coughing, fatigue and depressed mood are common during an acute infection with severe respiratory syndrome coronavirus 2 (SARS-CoV-2) or coronavirus disease 2019 (COVID). However, a substantial number of people report persistence of symptoms that last at least three months after infection with SARS-CoV-2, which has been defined by the World Health Organisation as POST-COVID syndrome (Soriano et al., 2022). Patients report a wide variety of physical and mood-related symptoms (Lopez-Leon et al., 2021), with fatigue and depression being the most commonly reported, but least understood, symptoms.

These symptoms considerably impact daily and occupational functioning. While fatigue is an adaptive feeling of physical or mental exhaustion that promotes rest-taking behaviour, depression is characterized by a reduced ability to experience pleasure (*Allostatic Self-Efficacy: A Metacognitive Theory of Dyshomeostasis-Induced Fatigue and Depression - PubMed*, n.d.; Dantzer et al., 2014). Both fatigue and depression affect daily functioning by changing decision making. Specifically, they have been associated with alterations in decisions to engage in activities that involve weighing of the costs (e.g. effort investments, or effort-sensitivity) against the benefits (e.g. the experience of pleasure, or reward-sensitivity) of that activity (Erfanian Abdoust et al., 2024; Matthews et al., 2023; Treadway et al., 2012, p. 202). However, how fatigue and depression relate to decision making and whether this is the same during acute and persistent phases of a COVID infection, is currently unclear. Assessing the behavioural consequences of fatigue and depression by means of decision making could therefore help to better understand the differential impact of these symptoms on daily and occupational functioning during acute and chronic phases of COVID infections.

During an acute microbial or viral infection, systemic inflammation induces a process known as sickness behaviour, which involves feelings of fatigue and depression, but also increases in rest-taking behaviours (Dantzer, 2001a). Sickness behaviour is considered to be an adaptive behavioural change that promotes recovery by directing bodily energy resources towards internal processes that fight disease and away from (effortful) physical or mental activities (Dantzer, 2001b). Studies in the field of cognitive neuroscience have investigated sickness behaviors using effort-based decision making tasks (Matthews et al., 2023; Müller & Apps, 2019; Treadway et al., 2012). In this type of decision making tasks, one decides whether or not to perform an effortful task by weighing the amount of effort it will cost against the reward it will bring (Chong et al., 2016). By presenting multiple offers with various combination of effort and reward levels, it is possible to analyze choice-behavior to derive individual parameters of effort- and reward-sensitivity, i.e. reflecting how effort and reward information influence decisions to engage in activities. Previous research using this type of tasks has shown that acute inflammation induced by bacterial lipopolysaccharide (LPS) in healthy individuals indeed alters effort-based decision making. Specifically, fatigue during acute systemic inflammation was selectively associated with increased sensitivity to effort but not reward information during decisions to engaging in a rewarding but effortful activity (Draper et al., 2018; Lambregts et al., 2023). These results provide insight into which processes are affected by acute inflammation to promote rest-taking behaviours during sickness.

When symptoms after acute infections persist, their characteristics often change, raising the question whether their relationship to decision making also changes. A study from Capuron & Miller (2011) showed that patients receiving long-lasting treatment with the immune-stimulant Interferon-alfa (IFN-alfa) initially only experienced vegetative sickness symptoms, such as fatigue and reduced appetite, in the first 4 weeks. However, after 4-6 weeks, they additionally started to develop mood and cognitive problems, including major depressive disorder (Capuron & Miller, 2011). Similar patterns were observed with COVID: Davis et al. (2021) showed that while fatigue (and other general sickness symptoms, e.g. fever) started directly after the infection, other symptoms, including cognitive problems and post-exertional malaise, i.e. worsening of symptoms following physical or mental exertion, started to increase after 4-8 weeks (Davis et al., 2021). This has led to the hypothesis that the mechanism through which inflammation causes acute versus long-term symptoms might be different. In line with that, depressive symptoms, which are also common in conditions of chronic low-grade inflammation (Beurel et al., 2020; Kohler et al., 2016; Miller et al., 2009) and reflect a reduced ability to experience pleasure, have been related to decreased sensitivity to reward (Keren et al., 2018; O’Callaghan & Stringaris, 2019; Treadway et al., 2012). Thus, while both fatigue and depressive mood result in similar changes in decision making, namely less engagement in effortful – but rewarding – activities, the mechanism driving this decision making might be different, and might change during the transition from an acute COVID infection to a post-COVID condition.

In this paper, we aimed to investigate how fatigue and depressive mood during different phases of a COVID infection relate to effort and reward sensitivity. We used a well-established computerized effort-based decision making task (Bonnelle et al., 2016; Contreras-Huerta et al., 2022) to test whether the relationships between effort/reward sensitivity and fatigue/depression symptoms differed between people who had COVID >12 weeks ago, individuals who had COVID <4 weeks ago and individuals who did not have COVID. We hypothesized that fatigue in both the <4 weeks and >12 weeks group is more strongly associated with increased effort sensitivity compared to the no COVID group. Additionally, we hypothesize that depressive mood is more strongly associated with decreased reward sensitivity only in the >12 weeks group compared to the <4 weeks and no COVID groups (Figure 1; preregistered at https://osf.io/uhwkc).

**Figure 1:**
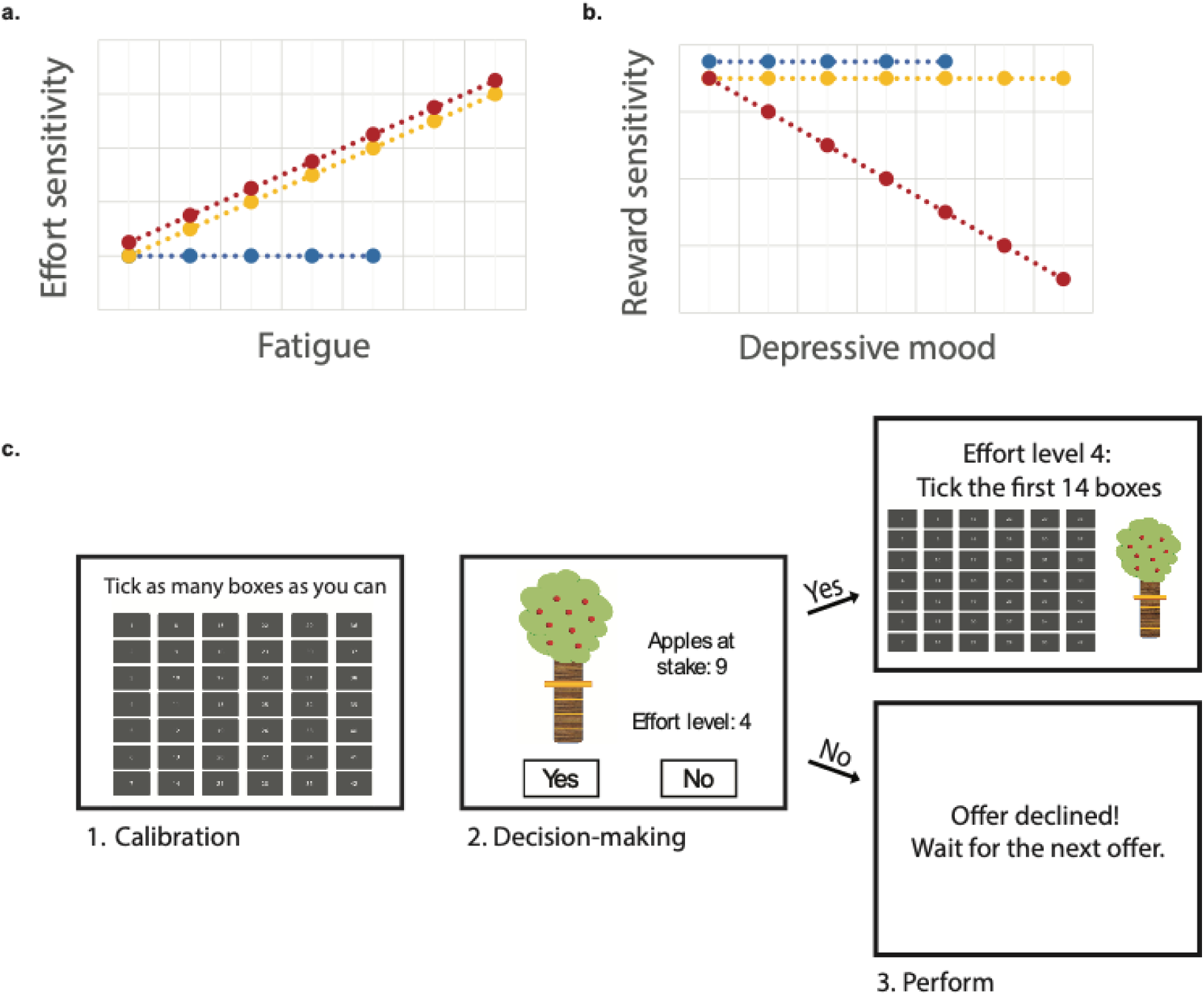
Visual overview of our hypothesis regarding the association between fatigue and effort sensitivity (a) and depressive mood and reward sensitivity (b), and the task design (c) of the study.

Currently, it is unclear why some people develop post-COVID and others do not. To better understand the transition from acute to persistent symptoms, it is important to understand who is at risk for developing post-COVID and possible associated alterations in decision making. Previous studies have already identified various factors that predict acute COVID severity, including lifestyle-related factors, such as BMI, smoking, age and sex (Chudzik et al., 2022; Kofahi et al., 2022; Merino et al., 2021; Pływaczewska-Jakubowska et al., 2022). For post-COVID these predictors are, however, less clear: some studies suggest that higher BMI, less exercise and smoking might also increase the risk for developing post-COVID (Bai et al., 2022; Pływaczewska-Jakubowska et al., 2022; Subramanian et al., 2022), while others do not find this association (Crook et al., 2021; Pływaczewska-Jakubowska et al., 2022). One reason for this heterogeneity, is that different studies used different ways of measuring post-COVID symptoms that included various combination of physiological, fatigue and mood-related symptoms, which largely vary between individuals. Accordingly, here we used the decision making task to investigate whether we can identify predictors of effort and reward sensitivity in people who had COVID >12 weeks ago, compared to individuals who had COVID <4 weeks ago, or did not have COVID. In previous cognitive neuroscience studies, similar lifestyle factors, such as BMI, age, and smoking have indeed been found to be associated with altered decision making (Addicott et al., 2020; Byrne & Ghaiumy Anaraky, 2020; Mansur et al., 2019). By focusing on the independent processes of decision making (i.e. effort and reward sensitivity), we might be able to delineate specific predictors for specific behavioral consequences of COVID.

Taken together, post-COVID symptoms of fatigue and depressive mood are currently poorly understood. This study uses effort-based decision making to investigate how fatigue and depressive mood during different phases of a COVID infection relate to effort and reward sensitivity, and assesses which risk factors predict effort and reward sensitivity in individuals who had COVID > 12 weeks ago. Results may provide insight into differential processes that underlie fatigue and depressive symptoms after COVID infections and how this impacts decisions about energy expenditure in daily functioning.

## Materials and methods

### Population and design

This online study had a cross-sectional design. We preregistered the study on OSF (available via https://osf.io/3vfku/). In total, we recruited 308 Dutch-speaking participants aged 18-65 years old through social media (e.g. post-COVID patient groups), flyers and the university’s online recruitment database. We recruited three groups of people: People who had COVID more than 12 weeks ago (n= 113), people who had COVID less than 4 weeks ago (n= 89) and people who never had COVID (self-reported) (n= 106). Recruitment took place between March 2021 and February 2022 (Supplementary Figure 1).

### Procedure

When opening the link of the study, participants first encountered a webpage with information about the study. Participants gave informed consent via an opt-in button, after which they were redirected to the study. Participants were then asked to fill in several questionnaires, and perform an online behavioural task, together taking approximately 30 minutes to complete. Among the participants, we distributed 40 gift cards of 5 euros plus up to 5 euros that participants were able to win during the behavioural task in a lottery draw. Participants performed the study in the online questionnaire platform Gorilla Experiment Builder (www.gorilla.sc) (*Gorilla in Our Midst: An Online Behavioral Experiment Builder | Behavior Research Methods*, n.d.; *Realistic Precision and Accuracy of Online Experiment Platforms, Web Browsers, and Devices | Behavior Research Methods*, n.d.; Tomczak et al., 2023), which could only be performed on a computer, laptop, or tablet (not on a smart phone).

### Measurement of fatigue and depression

We measured the main predictors in this study, current state fatigue (scale: 6-30) and depressive mood (scale: 8-40) (i.e. how do you feel right now), by fatigue and depressive mood subscales of the Profile of Mood States questionnaire (Grove & Prapavessis, 1992; McNair et al., 1971) before starting with the experimental task. In addition, for descriptive purposes, we additionally measured overall fatigue in the past two weeks by the Multidimensional Fatigue Inventory (MFI) (Smets et al., 1995).

### Experimental task: Effort-based decision making

We measured effort and reward sensitivity using an adjusted online version of the decision-making task from Bonelle et al. (2016) and Contreras-Huerta et al (2022) (Bonnelle et al., 2016; Contreras-Huerta et al., 2022) (Figure 1). In this task, participants decided whether a certain reward, i.e. apples in the tree, was worth a certain amount of effort, i.e. clicking a certain number of boxes within 10 seconds. The task was divided in four parts: Calibration, familiarization, decision-making and execution. The decision-making and execution phases were separated to avoid that fatigue due to the execution would affect the participant’s decisions.

During calibration, we measured how many boxes the participant was able to tick with their computer mouse or on their tablet’s touch screen within 10 seconds. This was repeated three times, and the highest number of boxes ticked (maximum number of boxes ticked (MBT)) was used to determine the five levels of effort (i.e. 10, 30, 50, 70, and 90% of MBT). During familiarization, participants performed all five levels of effort three times to get familiar with each effort level. To validate the experienced task load, participants filled in the NASA Task Load index after each effort level (Hart, 1986).

During the decision phase, we presented participants with offers of combinations consisting of one of 5 levels of effort (10, 30, 50, 70 and 90% of MBT) with one of 5 levels of reward (1, 3, 6, 9 and 12 apples). Participants had a maximum of ten seconds to decide whether they were willing to perform the amount of effort for the reward at stake. All 25 combinations of the effort and reward levels were presented to the participants five times, resulting in a total of 125 trials.

During the execution phase, participants executed 25 trials that were randomly selected from the decision phase. If they had accepted the trial, participants performed the trial and, if successful, collected the apples. Each collected apple corresponded to an additional €0,04 on the €5,-gift card they could win in a lottery. If they had not accepted the trial, participants waited for 10 seconds until the next trial was presented.

### Demographics and population characteristics

We collected information on demographics (including age, sex, education, socio economic status) and population characteristics using existing questionnaires and newly created items. We used the 36-item Short Form Health Survey (Hays et al., 1993) to describe the current physical health status of the population and the Multi-Dimensional Fatigue Inventory (MFI) to describe the fatigue levels in the past two weeks.

To assess symptoms associated with COVID infection and POST-COVID, we created a set of 29 of physical cognitive and mood symptoms (the 29 symptom list). We assess which symptoms participants experienced during the first two weeks of COVID infections on a 5-point scale (1=not at all, 2=Rarely, 3=Sometimes 4=Often, 5= Always) and we assessed the relative change in symptoms during the two weeks prior to participation as compared to before March 2020 on a 6-point scale (1=not at all, 2=much less often, 3=slightly less often 4=just as often, 5= slightly more often, 6=much more often). An overview of all items and questionnaires are shown in Supplementary Table 1.

### Predictors of effort and reward sensitivity

The items and questionnaires were used to compute the following predictors of effort and reward sensitivity: age (years), sex (male, female), pre-pandemic BMI (kg/m2), pre-pandemic lifestyle, socio-economic status, pre-pandemic physical health, COVID disease severity, worrying during the acute phase of COVID, and use of COVID-related aftercare.

For pre-pandemic lifestyle, socio-economic status, pre-pandemic physical health, and COVID disease severity, we calculated a combined score by averaging Z-scores of a set of related variables. Specifically, for lifestyle, we calculated a combined score by averaging Z-scores of pre-pandemic smoking (no, occasionally, or daily), alcohol use (never, monthly, 2-4 times a month, 2-3 times a week, or >4 times a week), medium and high intensive exercising (never, 1-2h a week, 2-3h a week, 4-5 a week, or >5h a week), and having diabetes type II, where a higher Z-score for lifestyle score indicates a healthier lifestyle.

For socio-economic status, we calculated a combined score by averaging Z-scores of income (€ per year) and educational level (ISCED score (*International Standard Classification of Education (ISCED)*, n.d.)).

For pre-pandemic physical health calculated a combined score by averaging Z-scores of the number of pre-pandemic chronic diseases participants had.

For COVID disease severity, we asked participants who had COVID to what extent they experienced a list of 29 symptoms during the first two weeks of their infection on a 5-point scale (1=not at all, 5=all the time). We then combined the number and severity of COVID symptoms, and type of medical care used because of COVID (none, GP, hospital admission, or intensive care admission) into one Z-score.

We quantified COVID-related worrying by asking to what extent people were worried about their own health during the first two weeks of their infection on a 5-point scale (1=never, 5=all the time). Use of COVID-related aftercare was defined by whether the participants received physical therapy, ergotherapy, social care or were referred to a neurologist, pulmonologist, psychologist, or psychiatrist after their COVID infection.

Finally, we measured pandemic-related stressor load with a questionnaire asking whether and to what extent participants experienced a list of 18 potential stressors on a 6-point scale (1=not bothersome, 6=very bothersome) (van der Heide et al., n.d.).

### Statistical analysis

#### Demographics

We report demographic participant characteristics as mean±SD or proportions. Differences between the three groups were tested with one-way ANOVAs and Tukey-HSD post-hoc tests for continuous variables, or chi-square tests for categorical variables.

#### Effort and reward sensitivity and fatigue/depressive mood

Following our preregistration, we studied the relationship between fatigue and depressive mood with effort and reward sensitivity using mixed binomial regression modelling. As an additional exploratory analysis, we also investigated the effort*reward interaction to assess how fatigue and depressive mood are related to the integration of effort and reward information during decision making.

First, to test whether the relationship between fatigue and effort, reward and its interaction differed between the groups, we constructed a mixed model with the decision to accept the trial’s offer or not (yes/no) as dependent variable, Effort, Reward and Effort*Reward as random factors, the interaction between Group, Fatigue, Effort and Reward as fixed factors of interest, and Age and Sex as fixed factors of no interest. Second, we tested whether the relationship with depressive mood and effort, reward and its interaction differed between the groups by replacing Fatigue with Depressive mood scores in the above-described model. We Z-scored all fixed and random variables in the models. For each model, estimates (betas) of Effort and Reward for each participant were derived for interpretation and visualization. These beta’s represent effort and reward sensitivity, i.e. the extent to which decisions to accept an offer are affected by effort and reward levels respectively. For similar interpretation of the betas for Effort and Reward level, we reversed the Effort levels (1= 90% MVC, 5= 10% MVC) in these regression models such that higher values represent lower effort levels. Now for both Effort and Reward more positive betas represent higher sensitivity.

### Prediction analysis

We tested the following nine predictors of effort and reward sensitivity: age (years), sex (male, female), pre-pandemic BMI (kg/m2), pre-pandemic lifestyle, socio-economic status, pre-pandemic physical health, COVID disease severity, worrying during the acute phase of COVID, and use of COVID-related aftercare. For the prediction analysis, we transformed all predictors to Z-scores. First, we tested whether these predictors were related to effort sensitivity, reward sensitivity and its interaction within each group by performing three mixed binomial regression models, one for each group, with the trial answers (yes/no) as dependent variable, Effort, Reward and Effort*Reward as random factors, and the interaction between the nine predictors, Effort level, and Reward level as fixed factors. Second, to test whether the relationship between the predictors and effort and reward sensitivity differed between the three groups, we added Group (<4wks, >12 wks, no COVID) as fixed factor to this model. Predictors that were only measured in one or two of the three groups were not included in this model (i.e. COVID disease severity, worrying during the acute phase of COVID, and use of COVID-related aftercare). For similar interpretation of the betas for Effort and Reward level, we again reversed the levels of Effort levels (1= 90% MVC, 5= 10% MVC) in these regression models such that higher values represent lower effort levels.

We considered p<0.05 significant. We performed regression analyses using RStudio (version 1.4, R Core Team (2021)), R Foundation for Statistical Computing, Vienna, Austria) using the packages lme4 and lmerTest (Bates et al., 2015; Kuznetsova et al., 2017).

## Results

### Population characteristics

In total, 233 out of the 308 participants who completed the online study were included in the study after pre-registered data quality control (https://osf.io/uhwkc), of which 90 never had COVID, 62 had COVID <4 weeks ago and 81 had COVID >12 weeks ago. 75 of the 308 participants were excluded because they 1) accepted more than 90% or less than 10% of the offers in the decision phase (n= 67), 2) they pressed only one button in >70% of the decision trials (n= 3), 3) they missed more than 10 decision trials (n= 1), their MBT was <10 (n= 3), 4) their mean reaction time was shorter than 500ms (n= 1), or 5) they were not successful in >50% of the trials in the execution phase (n= 0).

Table 1 shows the main demographic characteristics of the study population. The COVID >12 weeks groups reported significantly more fatigue (as measured with the MFI) and more limitations in daily functioning (as measured with the SF-36) compared to the COVID <4 weeks and no COVID groups (Table 1 and supplementary information), while no group differences were observed in emotional well-being or limitations due to emotional problems. The results from the 29 symptom list indicate that fatigue, fatigue after mild exertion, having a long recovery after exertion, concentration problems and a high heart rate were the most common symptoms across the three groups. They also show that the COVID <4 weeks and COVID >12 weeks groups reported overall more physical and fatigue-related symptoms compared to participants who did not have COVID, while symptoms of feeling depressed, having a low motivation, feeling stressed/anxious did not differ between the three groups. The three groups also experienced a similar amount of pandemic-related stress (Supplementary Table 2).

**Table 1:** Demographic, physical health, mental health, and COVID-related characteristics of the study population according to group.

|  | Total (n= 242) | No COVID (n= 90) | COVID <4 weeks (n= 62) | COVID >12 weeks (n= 81) |  |
| --- | --- | --- | --- | --- | --- |
|  | *Mean±SD or n (%)* | *Mean±SD or n (%)* | *Mean±SD or n (%)* | *Mean±SD or n (%)* | *P-value* |
| Age (years) | 35±13 | 28±10 | 36±13 | 43±12 | <0.001<sup>abc</sup> |
| Sex (% male) | 41 (17.7) | 21 (23.3) | 7 (11.4) | 13 (16.0) | 0.154 |
| BMI, pre-pandemic (kg/m<sup>2</sup>) | 24.3±4.7 | 22.3±2.9 | 23.7±3.6 | 27.0±5.8 | <0.001<sup>bc</sup> |
| Ethnicity (% Dutch) | 219 (94.0) | 81 (90.0) | 58 (93.5) | 80 (98.8) | 0.054 |
| Education (ISCED score) | 5.1±1.5 | 5.2±1.6 | 5.4±1.5 | 4.8±1.3 | 0.062 |
| Income (€) | 50,182±40,978 | 38,306±38,762 | 59,314±44,075 | 56,389±38,138 | 0.002<sup>ab</sup> |
| Smoking, pre-pandemic (% daily) | 8 (3.4) | 3 (3.3) | 2 (3.2) | 3 (3.7) | 0.465 |
| Alcohol use, pre-pandemic (times per month) | 4.3±5.1 | 3.7±4.6 | 5.7±6.0 | 3.8±4.7 | 0.035<sup>a</sup> |
| Medium intensive exercising, pre-pandemic (h per week) | 3.34±1.72 | 2.90±1.73 | 3.19±1.56 | 3.94±1.67 | 0.001<sup>bc</sup> |
| High intensive exercising, pre-pandemic (h per week) | 2.34±1.72 | 2.22±1.67 | 2.48±1.70 | 2.38±1.80 | 0.639 |
| Having chronic disease, pre-pandemic (%) | 49 (21) | 16 (18) | 6 (9.7) | 22 (27) | 0.028<sup>c</sup> |
| SF-36 (range 0-100) |  |  |  |  |  |
| Physical functioning | 73±28 | 91±13 | 74±24 | 51±27 | <0.001<sup>abc</sup> |
| Role limitations due to physical health | 43±45 | 77±34 | 33±41 | 14±32 | <0.001<sup>abc</sup> |
| Role limitations due to emotional problems | 58±44 | 57±42 | 59±46 | 58±44 | 0.970 |
| Energy/fatigue | 43±23 | 54±19 | 42±23 | 32±20 | <0.001 |
| Emotional well-being | 62±20 | 63±20 | 63±22 | 61±17 | 0.751 |
| Social functioning | 49±26 | 57±21 | 48±28 | 41±29 | <0.001<sup>b</sup> |
| Pain | 71±27 | 86±19 | 67±26 | 57±27 | <0.001<sup>abc</sup> |
| General health | 57±20 | 65±18 | 61±20 | 46±19 | <0.001<sup>bc</sup> |
| Multidimensional Fatigue Index |  |  |  |  |  |
| Total fatigue (range 0-100) | 66.9±19.4 | 57.4±17.0 | 67.9±19.8 | 76.7±16.6 | <0.001<sup>abc</sup> |
| General fatigue in past 2 weeks (range 0-20) | 14.8±4.4 | 12.4±4.0 | 15.1±4.0 | 17.3±3.7 | <0.001<sup>abc</sup> |
| Physical fatigue (range 0-20) | 13.9±5.1 | 11.4±4.5 | 14.0±4.7 | 16.7±4.5 | <0.001<sup>abc</sup> |
| Reduced motivation (range 0-20) | 11.2±4.0 | 10.0±3.6 | 11.7±4.8 | 12.0±3.6 | <0.002<sup>ab</sup> |
| Reduced activity (range 0-20) | 13.9±4.4 | 11.9±4.1 | 14.5±4.4 | 15.5±4.1 | <0.001<sup>ab</sup> |
| Mental fatigue (range 0-20) | 13.1±4.8 | 11.6±4.6 | 12.7±4.6 | 15.1±4.5 | <0.001<sup>bc</sup> |
| **COVID-related information** |  |  |  |  |  |
| COVID-related stressors (score, range 0-125) | 40±11 | 38±9 | 41±12 | 41±11 | 0.078 |
| Time since symptom onset (days) | 112±129 | N.a. | 15±27 | 190±125 | <0.001 |
| Symptom severity during first two weeks (score, range 0-89) | 45.3±22.4 | N.a. | 38.6±21.3 | 50.5±22 | 0.001 |
| Medical care used |  |  |  |  | <0.001 |
| *None* | 72 (50.7) | N.a. | 47 (75.8) | 25 (31.3) |  |
| *GP/ Emergency unit* | 63 (44.4) | N.a. | 15 (24.2) | 48 (60.0) |  |
| *Hospital admission* | 6 (4.2) | N.a. | 0 (0.0) | 7 (8.7) |  |
| Aftercare (% yes) | 66 (46.2) | N.a. | 6 (9.7) | 60 (74.1) | <0.001 |
| Vaccinated before COVID infection (% yes) | 36 (60.0) | N.a. | 34 (94.4) | 1 (4.3) | <0.001 |
| Not yet recovered at moment of participation (% yes) | 105 (73.9) | N.a. | 38 (61.3) | 67 (83.8) | 0.004 |
| Worrying about illness in first two weeks of infection (score, range 1-6) | 3.9±1.3 | N.a. | 3.8±1.4 | 4.0±1.2 | 0.419 |

BMI, body mass index; GP, general practitioner; ISCED, International Standard
Classification of Education; MFI, Multidimensional Fatigue Index; SF36, 36-Item Short Form Health Survey; POMS, Profile of Mood States.
P-values for differences between COVID groups were determined by performing a one-way ANOVA with Tukey-HSD post hoc test for continuous variables and a chi-square test for categorical variables.
<sup>a</sup>No COVID group differs significantly from the COVID <4 weeks group (P<0.05). <sup>b</sup>No COVID group differs significantly from the COVID >12 weeks group (P<0.05). <sup>c</sup>COVID <4 weeks group differs significantly from the COVID >12 weeks group (P<0.05).

The COVID >12 weeks group was older, had a higher pre-pandemic and current BMI, did more medium intensive exercising before the pandemic, and more often had a chronic disease compared to both the no COVID and COVID <4 weeks groups. The COVID >12 weeks group also had a higher income compared to the no COVID group and did more medium and high intensive exercising at the moment of participation compared to the COVID <4 weeks group.

Regarding their COVID infection, the COVID >12 weeks group was less often vaccinated before their infection, had a higher symptom severity, and were more worried about their illness in the first two weeks of their infection, received more often medical care, and indicated less often to be recovered at time of testing compared to the COVID <4 weeks group (Table 1). In addition, while most participants in the COVID <4 weeks group had COVID when the delta and omicron variants were predominant in the Netherlands, the COVID >12 weeks group had COVID when the alpha variant was predominant (Supplementary Figure 1).

### Fatigue and depressive mood at time of testing

The COVID >12 weeks group was more fatigued at time of testing compared to the COVID <4 weeks and no COVID groups (mean±SD: 20±7 vs. 14±7 and 12±6 POMS-score, respectively; both p<0.001) (Figure 2a). Depressive mood at time of testing did not differ between the groups (p=0.052) (Figure 2b).

**Figure 2:**
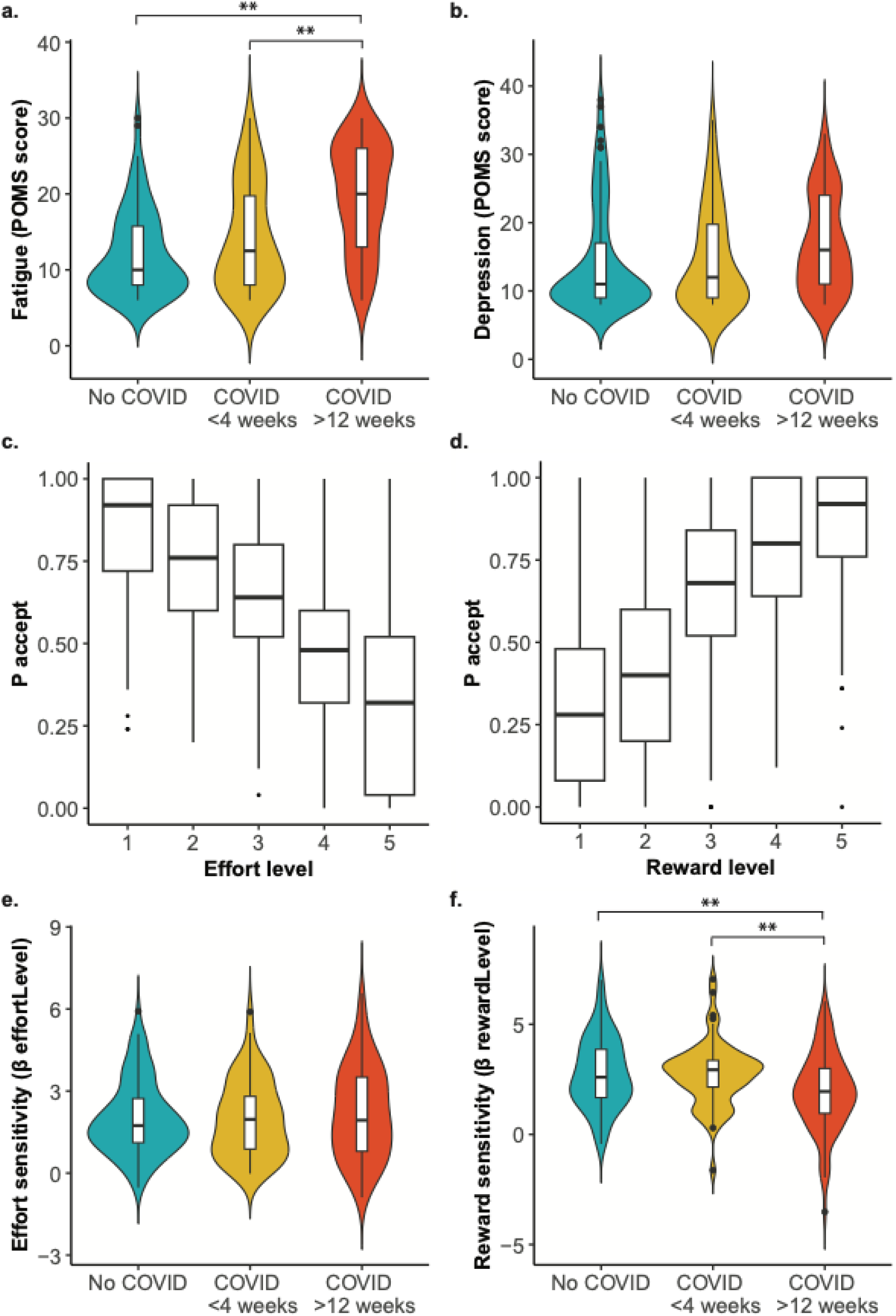
State fatigue (a) and depressive mood (b) scores at moment of participation according to group and mean acceptance rates for each effort (c) and reward (d) during the decision phase of the effort-based decision making task, and therefrom derived values for effort sensitivity (e), reward sensitivity (f) according to group. Regression coefficients for effort (e) and reward (f) were determined by mixed model binomial regression analysis. **P<0.01

### Effort-based decision-making task: Validation

In line with previous work using a similar task (Bonnelle et al., 2016), mixed binomial regression analysis showed a main effect of Effort (β 2.21 (95%CI: 1.98, 2.45); OR: 9.11 (95%CI: 7.24, 11.59), p<0.001) and Reward (β 2.66 (95% CI: 2.40, 2.93); OR: 14.30 (95%CI: 11.02, 18.73), p<0.001) across the three groups: Participants accepted more offers when the reward was high or when the required effort level was low (Figure 3c and 3d). The results of NASA task load Index were in line with this pattern: Participants experienced the increasing effort levels as more effortful (F=95.974, df=4, p<0.001), more physically (F=65.555, df=4, p<0.001), mentally (F=67.988, df=4, p<0.001) and time demanding (F=158.900, df=4, p<0.001), and frustrating (F=51.743, df=4, p<0.001), and these ratings did not differ between the groups (Supplementary Figure 4).

**Figure 3:**
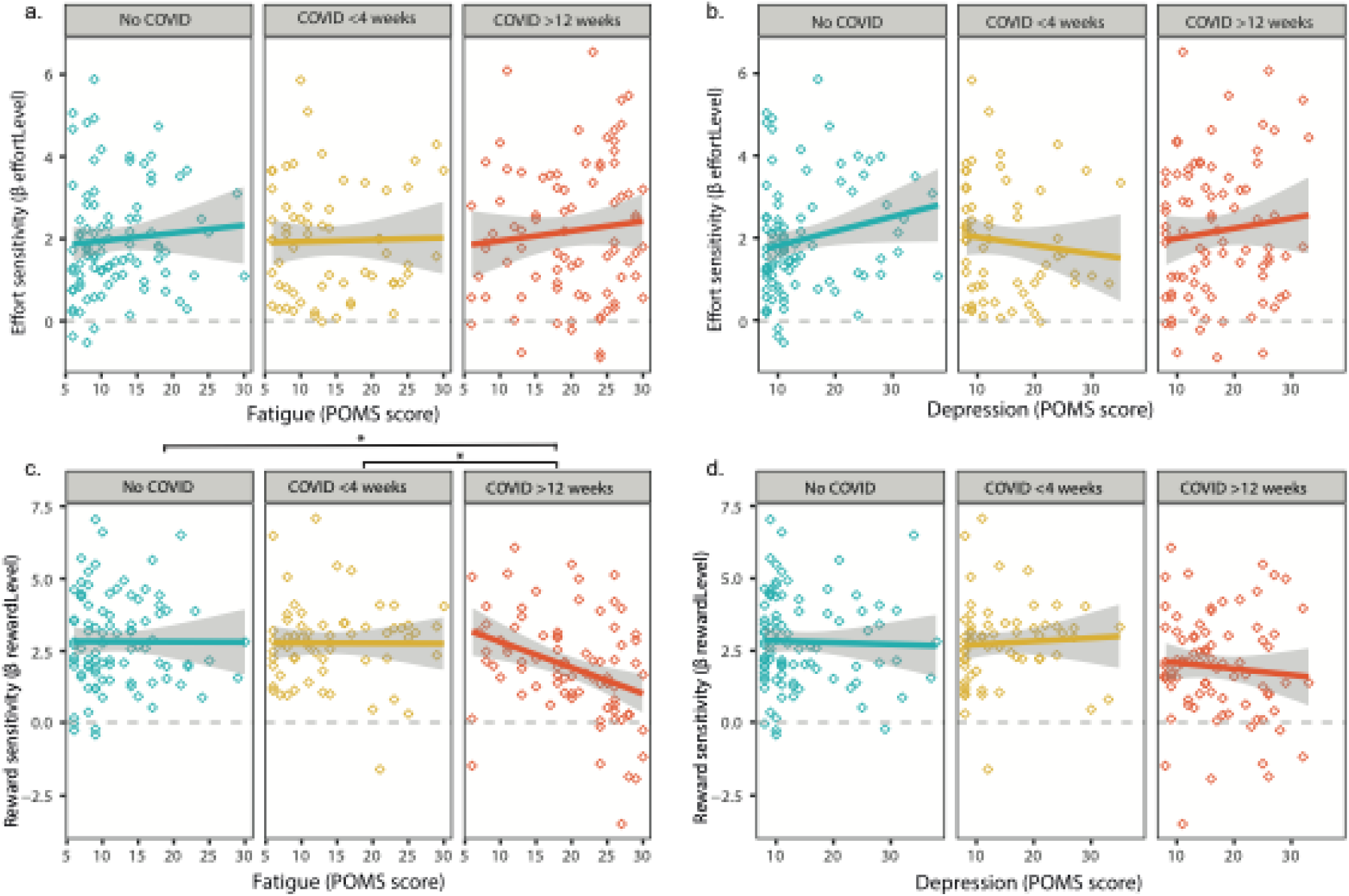
The association between fatigue and depressive mood and effort (a and b), and reward (c and d) according to group. The regression coefficients for effort and reward sensitivity, and the differences between groups were determined and tested by mixed model binomial regression analysis. A higher regression coefficient reflects more optimal decision making for effort or reward, i.e. higher effort or reward sensitivity. *P<0.05

The mean (standard deviation (SD)) maximum number of clicks during the calibration phase across all groups was 24±7 and did not differ between the groups (F=1.78, df=2, p=0.171), indicating that the groups did, on average, not differ in their ability to click the boxes. During the execution phase, the mean±SD success rate of the trials was 97±6%, and mean±SD total rewards obtained during the execution phase was 46±14 apples, corresponding with €5,-+ €1.84±0.56 worth in gift cards, all of which did not differ between the groups either (F=0.571, df=2, p=0.571), indicating that all participants were highly successful in executing the accepted offers.

### Effort-based decision-making task: Group differences

Reward sensitivity, the extent to which increasing reward levels affects the choice to accept the offer, were lower in the COVID >12 weeks group compared to the no COVID group (Reward*Group(COVID>12weeks vs. no COVID): β -1.05 (95% CI -1.63, -0.47) OR: 0.35 (95%CI 0.20, 0.62), p<0.001) and compared to the COVID <4 weeks group (Reward*Group(COVID>12weeks vs. COVID<4weeks): β -0.97 (95% CI -1.61, -0.34); OR: 0.38 (95%CI 0.20, 0.72), p=0.003) (Figure 2f).

Acceptance rates for effort sensitivity, i.e. the extent to which increasing effort levels affects the choice to accept the offer, did not differ between the groups (Figure 2e).

### Effort-based decision-making task: Relationship between Fatigue and effort and reward sensitivity

Mixed binomial regression analysis of decisions during the effort-based decision making task revealed interaction effects for Reward*Fatigue*Group: We found that the association between fatigue and reward sensitivity was more negative in the COVID >12 weeks group compared to the no COVID and the COVID <4 weeks group (β Reward*Fatigue*Group (COVID>12weeks vs. no COVID): -0.74 (95% CI -1.39, -0.08): OR 0.48 (95%CI 4.01, 0.92), p=0.029; β Reward*Fatigue*Group (COVID>12weeks vs. COVID<4weeks): -0.75 (95% CI -1.40, 0.12); OR: 0.47 (95%CI 0.25, 1.13), p=0.022, respectively). Visual inspection of the regression plots (Figure 3) suggests that, especially in high-fatigued individuals in the COVID >12 weeks group, the impact of reward information on decisions is lower compared to the NO-COVID and the COVID <4 weeks group, while those reporting low fatigue showed reward sensitivity levels comparable to the other groups.

We did not observe interactions between Effort*Fatigue*Group, indicating that the relationship between fatigue and effort sensitivity did not differ between groups (all β <0.39, all OR <1.47, all p>0.05). We also did not observe an Effort*Fatigue interaction, indicating that, across the three groups, fatigue was not associated with effort sensitivity (β Effort*Fatigue: 0.15 (95% CI -0.32, 0.63): OR 1.16 (95%CI 0.73, 1.87), p=0.520).

### Effort-based decision-making task: Relationship between depressive mood and effort and reward sensitivity

We did not observe any interactions between Reward*DepressiveMood*Group or between Effort*DepressiveMood*Group, meaning that the relationship between depressive mood and reward or effort sensitivity did not differ between the groups (all β <0.49, all OR <1.63, all p>0.05). We also did not observe an Reward*DepressiveMood or Effort*DepressiveMood interaction, indicating that, across the three groups, DepressiveMood was not associated with reward or effort sensitivity (β Effort*DepressiveMood: 0.32 (95% CI -0.03, 0.66): OR 1.37 (95%CI 0.97, 1.94), p=0.075; Reward*DepressiveMood: -0.04 (95% CI -0.43, 0.66): OR 0.96 (95%CI 0.65, 1.42), p=0.841).

The results regarding the relationship between fatigue and depressive mood and the interaction of effort and reward can be found in Supplementary Figure 6.

### Effort-based decision-making task: Predictors of effort and reward sensitivity in the COVID>12 weeks group

For our secondary aim to identify risk factors of effort and reward sensitivity, we focused on the relationships between predictors and effort or reward sensitivity within the COVID >12 weeks group and how they differed from the other two groups.. All predictors of effort and reward sensitivity and their coefficient estimates are shown in Figure 4. The correlation matrix of the different predictors across groups can be found in Supplementary Figure 3.

**Figure 4:**
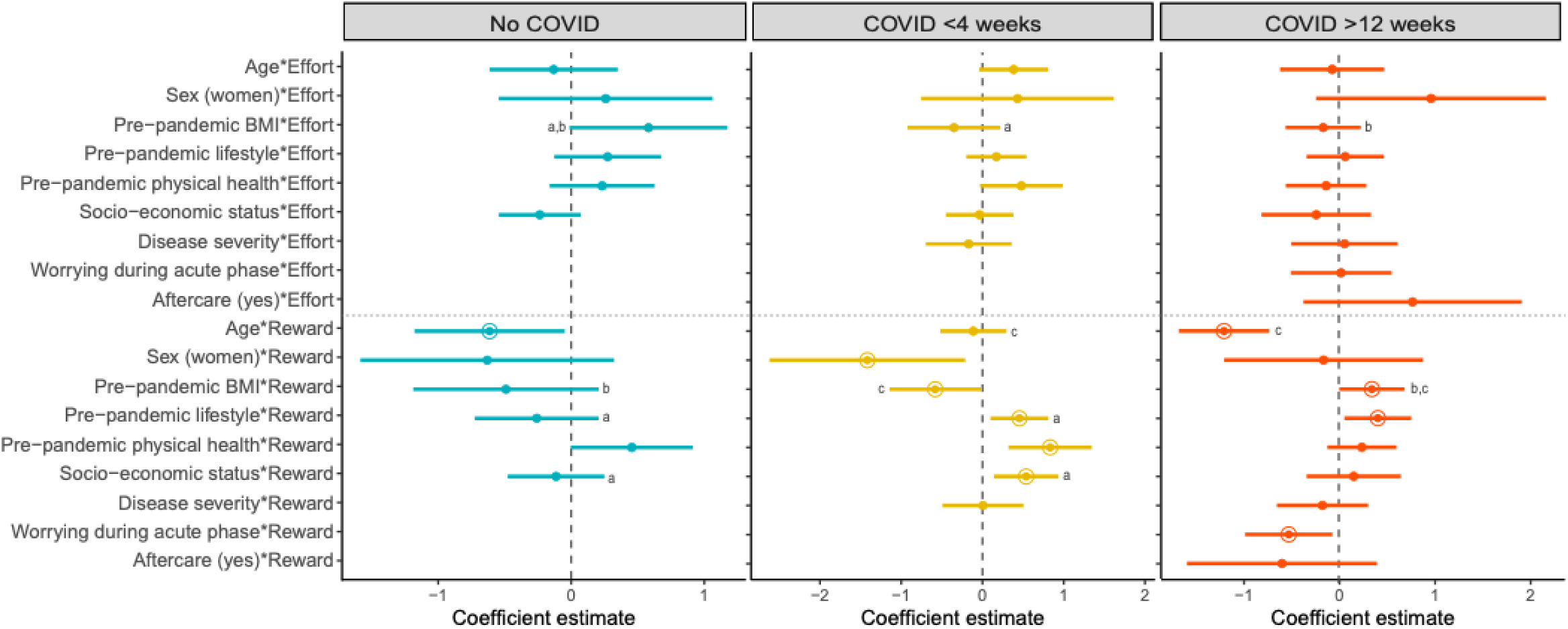
Prediction regression models of effort and reward in each group. Regression coefficients were estimated using mixed model binomial regression analysis. Continuous predictors were Z-scored. Error bars represent the 95% confidence interval. Significant regression coefficients are highlighted by a circle. A higher regression coefficient for effort or reward reflects more optimal decision making for effort or reward, i.e. higher effort or reward sensitivity. ^a^No COVID group differs significantly from the COVID <4 weeks group (P<0.05). ^b^No COVID group differs significantly from the COVID >12 weeks group (P<0.05). ^c^COVID <4 weeks group differs significantly from the COVID >12 weeks group (P<0.05).

Regarding reward sensitivity, we found that in the COVID >12 weeks group, higher age and worrying during the acute phase of COVID were associated with lower reward sensitivity while a higher pre-pandemic BMI and a healthier pre-pandemic lifestyle were associated with higher reward sensitivity (β Age*Reward: -1.20 (95%CI - 1.68, -0.73); OR 0.30 (95%CI 0.19, 0.48), p<0.001); β Worrying*Reward: -0.52 (95%CI -0.98, -0.07); OR 0.59 (95%CI 0.38, 0.94), p=0.025; β BMI*Reward: 0.35 (95%CI 0.01, 0.69); OR 1.43 (95%CI 1.01, 2.00), p=0.047; β Lifestyle*Reward: 0.41 (95%CI 0.06, 0.76); OR 1.50 (95%CI 1.06, 2.14), p=0.022)) (Supplementary figure 7b, 7,d, 7e, and 7h), When comparing the groups, we found that in the COVID >12 weeks group, age was more negatively associated with reward sensitivity compared to the COVID <4 weeks group (β Age*Reward*Group(COVID >12 weeks vs. COVID <4 weeks): 0.86 (95% CI 0.13, 1.58); OR 2.36 (95%CI 1.14, 4.88), p=0.020). In addition, in the COVID >12 weeks group, BMI was more positively associated with reward sensitivity compared to the COVID <4 weeks group (β Age*Reward*Group(COVID >12 weeks vs. COVID <4 weeks): -0.94 (95% CI -1.61, -0.27); OR 0.39 (95%CI 0.20, 0.76), p=0.006. No differences between the COVID>12 weeks group and the other two groups were observed for the relationships and the other predictors and reward sensitivity (all p >.05).

Regarding effort sensitivity, we did not find significant predictors of effort sensitivity within the COVID >12 weeks group. However, when comparing the groups, we found that, pre-pandemic BMI was more negatively associated with effort sensitivity in the COVID >12 weeks group, compared to the no COVID group (β BMI*Effort*Group(COVID >12 weeks vs. no COVID): -0.73 (95%CI -1.45, -0.01); OR 0.48 (95%CI 0.23, 0.99), p=0.048)) (supplementary figure 7a).

The results regarding the interaction of effort and reward can be found in Supplementary Figure 6. Regression plots of the significant associations between the predictors and effort or reward sensitivity for each group are shown in Supplementary Figure 7.

## Discussion

People with persisting symptoms after COVID commonly report fatigue and depressive mood as primary symptoms, but is unclear how these symptoms relate to decisions regarding effortful and rewarding activities, and whether this differs from fatigue and depression experienced during the acute COVID phase. Here, we investigated how self-reported fatigue and depressive mood during task performance were associated with different components of decision making (i.e. effort and reward sensitivity) in three groups: A group who had COVID >12 weeks ago, a group who had COVID <4 weeks ago and a group who did not have COVID. In addition, we aimed to identify which risk factors were associated with effort and reward sensitivity in the > 12 weeks group. Results indicated that higher self-reported fatigue >12 weeks after COVID was associated with lower reward, but not effort, sensitivity during decision making, while depression was not associated with either of these decision-making parameters. In addition, higher age, a less healthy lifestyle, a lower pre-pandemic BMI and more worrying during the acute phase of COVID were predictors of lower reward sensitivity in the COVID >12 weeks group. These results provide valuable insights into how persisting fatigue after COVID may impact decision making in daily functioning.

Participants in the > 12 weeks group showed on average lower reward sensitivity during decision making than the >4 weeks or no COVID groups and this reduction in reward sensitivity was most prominent when fatigue was high. This might suggest that persisting fatigue >12 weeks after COVID may involve problems in reward processing. In line with this, several lines of research suggest that post-COVID may involve persistent (neuro) inflammation, indicated by increased levels of pro-inflammatory cytokines such as TNF-α, IL-1β, and IL-17A (Ceban et al., 2022; Monje & Iwasaki, 2022) can affect cortical and subcortical brain networks that are involved in motivation and reward processing (Guedj et al., 2021; Sollini et al., 2021). Additionally, several studies have suggested that chronic, but not acute, inflammation can induce long-lasting changes in neural reward processing (Capuron et al., 2012; Felger & Lotrich, 2013), potentially through a reduction in the availability of dopamine receptors and impaired release of dopamine, thereby diminishing reward sensitivity. Indeed, a study from Lamontagne et al. (2021) observed anhedonia or a loss of interest in previously rewarding activities in post-COVID patients compared to healthy controls. Accordingly, these studies suggest that the observed low reward sensitivity in high-fatigued individuals >12 weeks after COVID may reflect alterations in neural reward processing resulting from long-term effects of (neuro) inflammation. To further confirm this suggestion, future neuroimaging studies could further investigate how altered decision making in POST-COVID conditions is related to neuro-inflammatory processes and dopamine network function.

Reward sensitivity was, in contrast to our hypothesis, not associated with depressive mood, which also involves anhedonia and loss of interest in rewarding activities. Nor did the relationship between depressive mood and reward sensitivity differ between the groups. This might be due to the observation that depressed mood, both during task performance and in the 2 weeks prior to participation, did not differ between the three groups. Additionally, all groups also reported a similar increase in depressive symptoms compared to before the COVID pandemic. Accordingly, this increase in depressive symptoms across all groups might suggest that depression symptoms may not result from the COVID infection, but may instead reflect a general response to pandemic-related societal changes, such as lockdowns and the overall stress associated with the global health crisis (COVID Mental Disorders Collaborators, 2021; Wu et al., 2021). Previous studies of the symptom profile of COVID also reported that while depression is a prevalent characteristic of post-COVID in 12% of the cases, fatigue was more prevalent, occurring in 58% of the cases (Lopez-Leon et al., 2021). Therefore, it is possible that fatigue in the >12 weeks group is a primary consequence of the infection, and may therefore more readily reflect infection-related effects on neural reward processing, while depressive mood may reflect the general emotional response to the pandemic-related restrictions that may not involve infection-related effects on neural reward processing.

Fatigue was, in contrast to our hypothesis, not related to effort sensitivity. Nor did the relationship between fatigue and effort sensitivity differ between the groups. Accordingly, the results in the <4 weeks and >12 weeks group did not mirror those seen after well-controlled acute immune manipulations, using LPS (Draper et al., 2018; Lambregts et al., 2023), which show that fatigue during 1-3 hours after LPS administration is associated with increased effort sensitivity. It might be possible that this relationship with effort sensitivity may only surface during severe acute illness and that the level of sickness in the <4 weeks group was too variable with some already recovered, while others may still feel sick, or that participants may not have performed the task if they felt too ill. Given that this was an online experiment, we could not control this. Future studies could assess decision making shorter after initial infection and assess inflammatory markers such as CRP to make sure that the task is assessed during the acute phase of sickness.

Other studies have shown that momentary fluctuations in state fatigue during task performance in healthy individuals was associated with effort sensitivity during subsequent decisions to engage in the task or rest, (Matthews et al., 2023; Müller & Apps, 2019). This suggests that fatigue can also be associated with effort sensitivity irrespective of disease status. We also did not find a relationship between fatigue and effort sensitivity across the groups. It might be possible that the variation in the study population which included healthy individuals, individuals who were acutely ill, who were recovered form COVID and who developed POST-COVID syndrome may have occluded this effect of state fatigue. Furthermore, in the current study, we separated the decision phase from the execution phase to rule out potential fatigue fluctuations due to task execution, which makes it less comparable to these studies in healthy individuals. Nevertheless, although effort sensitivity did not differentiate between the groups, the findings with respect to reward sensitivity still indicate that the fatigue experienced in the >12 weeks phase, as compared to fatigue in the acute phase, may be driven by different biological mechanisms, possibly altered dopamine transmission due to an extended period of raised inflammation levels.

We identified several risk and protective factors of reward sensitivity in this study. Higher age was associated with decreased reward msensitivity in the COVID >12 weeks and the no COVID groups. This is in line with previous neurocognitive studies, in which age was associated with lower reward sensitivity irrespective of infection status (Dhingra et al., 2020; Eppinger et al., 2012; Kardos et al., 2017). With age, the brain becomes more vulnerable to the inflammatory and neurological effects of COVID (Keller et al., 2022), possibly explaining the stronger link between age and reward sensitivity in the COVID >12 weeks group compared to the other groups.

Furthermore, we found that a healthier lifestyle was protective against reward deficits in people who had COVID >12 weeks, which is in line with studies showing that unhealthy lifestyle factors, e.g. low exercise and smoking, are related to higher symptom severity of acute COVID and high risk of post-COVID (Bai et al., 2022; Kofahi et al., 2022; Patanavanich & Glantz, 2020; Pływaczewska-Jakubowska et al., 2022; Subramanian et al., 2022). Lifestyle factors like smoking and low exercise have been associated with low-grade inflammation (Teixeira de Lemos et al., 2011; Walsh et al., 2011; Yanbaeva et al., 2007), possibly contributing to the inflammatory effects of (post) COVID on the brain’s reward networks.

Notably, while a higher BMI was related to increased sensitivity to rewards in the COVID >12 weeks group, it was associated with decreased sensitivity to rewards in the COVID <4 weeks group. This last finding is in line with COVID studies consistently finding BMI as a predictor of acute COVID severity (Kofahi et al., 2022; Subramanian et al., 2022; Vassilopoulou et al., 2022; Zhou et al., 2021), while for post-COVID this link is less consistent (Bai et al., 2022; Townsend et al., 2020). In healthy populations, higher BMI is linked to increased sensitivity to rewards (Miras et al., 2012; Sutton et al., 2022; Val-Laillet et al., 2015), which is in line with the finding in the COVID >12 weeks group. On the other hand, obesity has been linked to higher inflammation levels and worse symptom severity of acute COVID (Kofahi et al., 2022; Subramanian et al., 2022; Vassilopoulou et al., 2022; Zhou et al., 2021), fitting with our results in the COVID <4 weeks group. We therefore hypothesize that post-COVID-related inflammation is lower than acute COVID-related inflammation in people with a higher BMI. In acute COVID the effect of BMI might be overruled by the larger inflammatory response, decreasing reward sensitivity. Future studies are needed to confirm this hypothesis about the role of BMI in COVID.

Finally, worrying during the acute phase of COVID was associated with less sensitivity to rewards. Worrying during COVID may reflect stress-related processes, which is in line with the finding that anxiety contributes to post-COVID fatigue (Calabria et al., 2022). These findings show that individuals with a higher age, unhealthy lifestyle, and those who worry more during the first phase of COVID might be at risk of post-COVID-related symptoms, which underscores the importance of a multifaceted approach in understanding and treating post-COVID-related symptoms.

This study had several limitations. First, the population in the >12 weeks group was a mix of recovered individuals and individuals with a diagnosis of POST-COVID and we do not know whether they had a diagnosis for POST-COVID. We can therefore not conclude that the reduction in reward sensitivity is a characteristic of POST-COVID syndrome or merely fatigue after COVID. However, there are several factors indicating that part of our population was dealing with (post) COVID symptomatology. Participants in the >12 weeks group were recruited from social media support groups for post-COVID with the goal to bias recruitment to include more participants with post-COVID-related fatigue. As expected with this recruitment strategy, participants in the COVID >12 weeks group were not only more fatigued at the moment of participation, they also reported to be more fatigued in the two weeks prior to participation compared to the other two groups, followed by participants in the COVID <4 weeks group, as measured by the MFI. In addition, more than 80% of the participants who had COVID >12 weeks ago reported that they were not yet recovered from their COVID infection, thereby meeting the WHO-definition of post-COVID (Soriano et al., 2022). It is therefore likely that our results may relate to the proposed neurobiological mechanisms of POST-COVID syndrome (REF Peluso 2024 mechanisms of POST COVID). To further confirm this, future studies could assess decision making in a properly diagnosed population and compare it to other chronic diseases that involve fatigue and healthy individuals.

Second, our recruitment method also resulted in a selection bias, as seen in demographical differences across groups. Participants in the COVID >12 weeks group were older, had a higher BMI, and had more chronic diseases compared to the other groups, and experienced more severe symptoms during acute COVID. These group differences could have affected the predictor analysis, creating more variability in the >12 weeks group and these results should therefore be interpreted with caution. Future studies should carefully control and match the different groups on these demographic characteristics.

Third, we used a measure of state fatigue (i.e. fatigue at the moment of testing) rather than trait fatigue (i.e. fatigue in the past two weeks) in our analysis. As in previous studies, we used state fatigue as this relates better to current task performance compared to trait fatigue (van der Schaaf et al., 2018). Moreover, not all participants in the COVID <4 weeks group had COVID for more than 2 weeks, meaning that the trait fatigue score (which measured across the past two weeks) could not be used for them.

In conclusion, we found evidence that fatigue, but not depressive mood, occurring >12 weeks after COVID infection is associated with reward deficits compared to people in the acute phase of COVID or who did not have COVID. In addition, higher age, unhealthy lifestyle, and worrying during the acute phase of COVID are potential risk factors for developing reward deficits after >12 weeks. These results indicate that assessment of decision making can provide valuable insights into the characteristics of COVID-related fatigue and depression which may ultimately lead to new treatment targets. For example, given our finding that symptoms of fatigue in long-duration COVID were associated with reduced reward sensitivity, treatments targeting the neural reward system might provide valuable alternatives or additions to current treatment strategies. One avenue might for example be pharmacological stimulation of dopamine transmission with agonists or reuptake inhibitors. To progress towards such treatments, future studies should further investigate potential neuroimmune processes that affect reward processing after COVID infections.

## Abbreviations

ANOVA: Analysis of variance
COVID: Coronavirus disease 2019
BMI: Body mass index
IFN: Interferon
IQR: Interquartile range
ISCED: International Standard Classification of Education
MBT: Maximum number of boxes ticked
MFI: Multidimenstional Fatigue Index
POMS: Profile of Mood States
SF-26: 36-Item Short Form Health Survey
95% CI: 95% confidence interval

