## Supplementary materials for "Fatigue >12 weeks after coronavirus disease (COVID) is associated with reduced reward sensitivity during effort-based decision making"

[#]These authors contributed equally

**SUPPLEMENT**

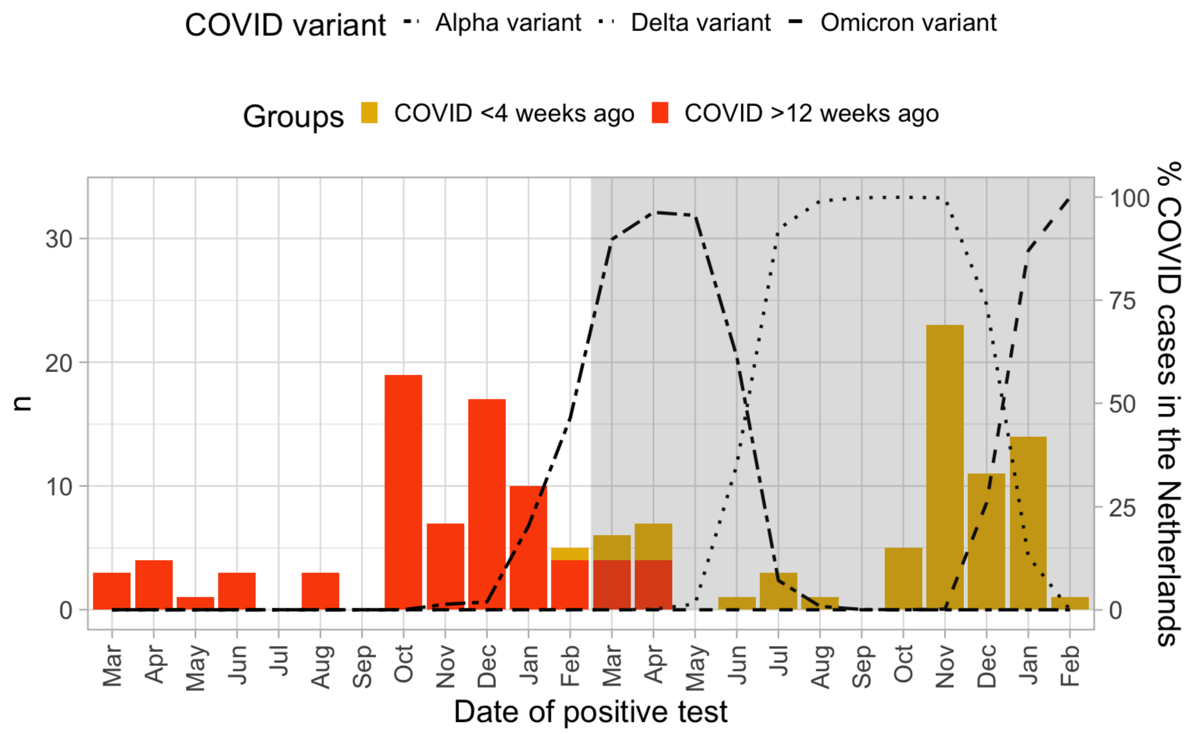


Supplementary Figure 1: Overview of inclusions of participants who had COVID <4 weeks and >12 weeks ago, and active COVID variants at that moment in the Netherlands (*COVID-19* | *RIVM*, n.d.).

Supplementary Table 1: Pre-pandemic and current health and lifestyle characteristics of the study population according to group.

| | Pre-pandemic | | | | Current | | | | Change |
|---|---|---|---|---|---|---|---|---|---|
| | **No COVID (n=90)** | **COVID <4 weeks (n=62)** | **COVID >12 weeks (n=81)** | | **No COVID (n=90)** | **COVID <4 weeks (n=62)** | **COVID >12 weeks (n=81)** | | **No COVID (n=90)** |
| | ***Mean±SD or n (%)*** | ***Mean±SD or n (%)*** | ***Mean±SD or n (%)*** | ***P-value*** | ***Mean±SD or n (%)*** | ***Mean±SD or n (%)*** | ***Mean±SD or n (%)*** | ***P-value*** | ***Mean±SD or n (%)*** |
| BMI (kg/m2) | 24.3±4.7 | 22.3±2.9 | 27.0±5.8 | <0.001[bc] | 24.6±4.9 | 22.6±3.0 | 27.5±5.9 | <0.001[bc] | 0.35±1.03 |
| Smoking (% daily) | 3 (3.3) | 2 (3.2) | 3 (3.7) | 0.465 | 2 (2.2) | 2 (3.2) | 1 (1.2) | 0.938 | -1 (-1.1) |
| Alcohol use (times per month) | 3.7±4.6 | 5.7±6.0 | 3.8±4.7 | 0.035[a] | 3.2±4.4 | 3.4±4.0 | 2.1±3.7 | 0.111 | -0.5±3.8 |
| Medium intensive exercising (h per week) | 2.90±1.73 | 3.19±1.56 | 3.94±1.67 | 0.001[bc] | 2.70±1.65 | 1.39±1.39 | 2.31±1.64 | <0.001[ac] | -0.20±2.08 |
| High intensive exercising (h per week) | 2.22±1.67 | 2.48±1.70 | 2.38±1.80 | 0.639 | 1.46±1.48 | 0.55±1.11 | 1.10±1.47 | <0.001[ac] | -0.77±1.65 |
| Having chronic disease (%) | 16 (18) | 6 (9.7) | 22 (27) | 0.028[c] | 16 (18) | 7 (11) | 26 (32) | 0.006[bc] | 0 (0.0) |

BMI, body mass index.

P-values for differences between COVID groups were determined by performing a one-way ANOVA with Tukey-HSD post hoc test for continuous variables and a chi-square test for categorical variables.

[a]No COVID group differs significantly from the COVID <4 weeks group (P<0.05). [b]No COVID group differs significantly from the COVID >12 weeks group (P<0.05). [c]COVID <4 weeks group differs significantly from the COVID >12 weeks group (P<0.05).

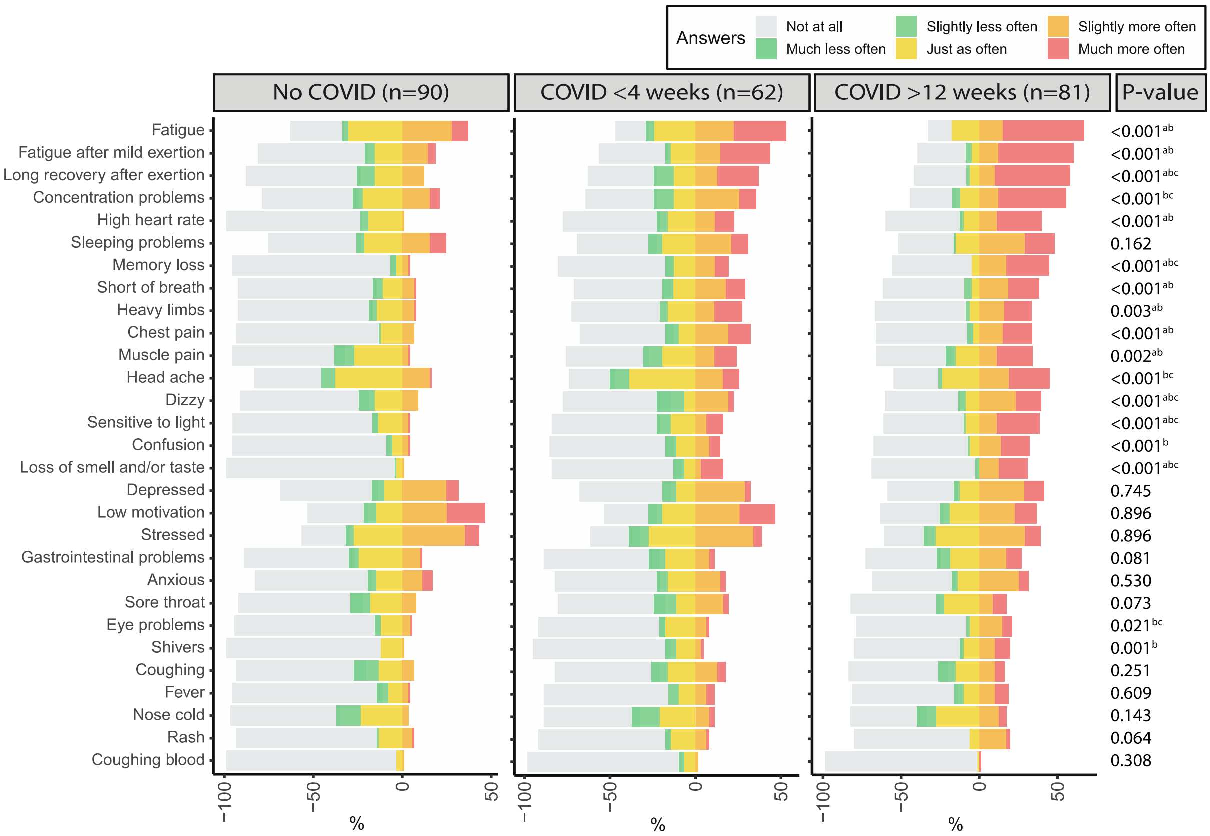


Supplementary Figure 2: Physical and psychological symptoms at the moment of participation compared to before the pandemic.

P-values for differences between groups were determined by performing chi-square tests.

[a]No COVID group differs significantly from the COVID <4 weeks group (P<0.05). [b]No COVID group differs significantly from the COVID >12 weeks group (P<0.05). [c]COVID <4 weeks group differs significantly from the COVID >12 weeks group (P<0.05).

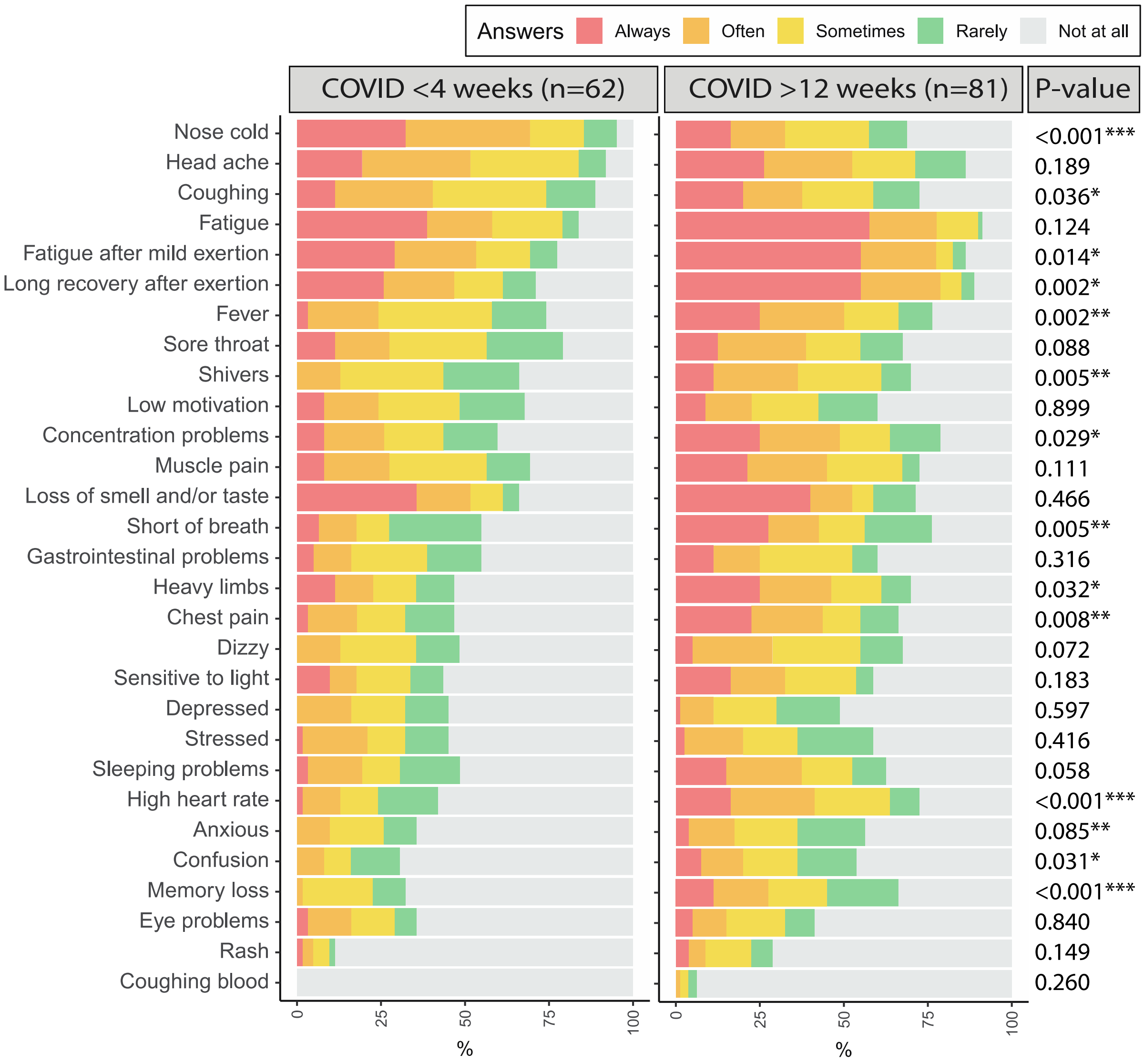


Supplementary Figure 3: Physical and psychological symptoms in the first 2 weeks of the acute infection compared to before the pandemic.

P-values for differences between groups were determined by performing chi-square tests.

*P<0.05, **P<0.01, ***P<0.001

Supplementary Table 2: COVID-related stressor load according to group.

| Stressors | No COVID (n=90) | | Acute COVID (n=62) | | Long COVID (n=81) | | P-value frequency | P-value burden |
|---|---|---|---|---|---|---|---|---|
| | Frequency | Average burden (1-5) | Frequency (%) | Average burden (1-5) | Frequency (%) | Average burden (1-5) | | |
| | n (%) | Mean±SD | n (%) | Mean±SD | n (%) | Mean±SD | | |
| Loss of social contact and social events | 86 (96) | 2.55±0.87 | 60 (97) | 2.74±1.05 | 76 (94) | 2.65±1.07 | 0.796 | 0.616 |
| COVID-19 related media coverage | 76 (84) | 2.83±1.42 | 57 (92) | 3.35±1.38 | 76 (94) | 3.20±1.36 | 0.104 | 0.083 |
| Family, friends, or loved ones working in vital professions | 53 (59) | 2.74±1.75 | 38 (61) | 2.82±1.76 | 56 (69) | 2.73±1.68 | 0.361 | 0.954 |
| Feeling restricted to leave your home | 52 (58) | 2.23±1.42 | 56 (90) | 2.70±1.14 | 57 (70) | 2.76±1.65 | <0.001[ac] | 0.053 |
| Family, friends, or loved ones being at increased risk for a serious course of the disease in case of a COVID-19 infection | 59 (66) | 2.21±1.33 | 46 (74) | 2.38±1.33 | 58 (72) | 2.33±1.35 | 0.481 | 0.765 |
| Having COVID-19 symptoms, or symptoms that could be related | 33 (37) | 1.87±1.47 | 59 (95) | 3.00±1.05 | 69 (85) | 2.60±0.93 | <0.001[ab] | <0.001[ab] |
| Being at risk for an infection (e.g., at work, in the supermarket) | 54 (60) | 2.36±1.53 | 45 (73) | 2.68±1.52 | 62 (77) | 2.63±1.45 | 0.051 | 0.414 |
| Not able to perform physical activity as usual | 65 (72) | 2.37±1.41 | 54 (87) | 2.55±1.18 | 71 (88) | 2.67±1.10 | 0.014[ab] | 0.393 |
| COVID-19 symptoms, or symptoms that could be related, in family members, friends, loved ones or colleagues | 50 (56) | 2.15±1.48 | 46 (74) | 2.44±1.36 | 48 (59) | 2.20±1.38 | 0.057 | 0.478 |
| Conflicts or disagreements in the family, social or professional environment | 41 (47) | 1.80±1.15 | 40 (65) | 1.93±1.09 | 49 (61) | 2.00±1.21 | 0.053 | 0.560 |
| Increased workload or work-related obstacles | 36 (40) | 1.58±0.92 | 36 (58) | 2.15±1.39 | 46 (57) | 1.81±1.12 | 0.036[ab] | 0.017[a] |
| Family, friends or loved ones are at the hospital and you are restricted in visiting them | 47 (52) | 1.82±1.25 | 28 (45) | 1.75±1.22 | 42 (52) | 1.90±1.22 | 0.649 | 0.802 |
| Severe disease or psychiatric problems of yourself or a loved one | 37 (42) | 1.68±1.08 | 23 (37) | 1.49±0.93 | 43 (54) | 1.91±1.29 | 0.114 | 0.125 |
| Tensions at home or family conflict | 33 (37) | 1.63±1.11 | 32 (52) | 2.04±1.39 | 37 (46) | 1.90±1.26 | 0.173 | 0.141 |
| Problems with access to healthcare, medication, or sanitation | 32 (36) | 1.80±1.38 | 24 (39) | 1.73±1.23 | 42 (52) | 2.23±1.56 | 0.081 | 0.075 |
| Being at increased risk for a serious course of the disease in case of an infection (belonging to a risk group) | 23 (26) | 1.60±1.28 | 24 (39) | 1.86±1.46 | 36 (44) | 1.95±1.39 | 0.030[b] | 0.258 |
| Problems obtaining basic needs and services | 20 (22) | 1.40±0.99 | 31 (50) | 2.12±1.50 | 30 (37) | 1.80±1.33 | 0.002[ab] | 0.003[a] |
| Financial problems | 28 (32) | 1.62±1.20 | 16 (26) | 1.52±1.11 | 22 (28) | 1.59±1.25 | 0.695 | 0.871 |
| Unable to attend a funeral of a loved one | 17 (19) | 1.28±0.81 | 15 (24) | 1.45±1.00 | 22 (27) | 1.42±0.85 | 0.430 | 0.462 |
| (Threat of) job loss, insolvency of a private company, for yourself or someone in your household | 13 (14) | 1.13±0.43 | 8 (13) | 1.28±0.88 | 19 (23) | 1.36±0.90 | 0.172 | 0.133 |

| | | | | | | | | |
|---|---|---|---|---|---|---|---|---|
| Death of a loved one | 16 (18) | 1.26±0.73 | 17 (27) | 1.27±0.62 | 10 (13) | 1.20±0.63 | 0.077 | 0.792 |
| Difficulties combining work with childcare | 6 (6.7) | 1.10±0.45 | 9 (15) | 1.25±0.77 | 12 (15) | 1.16±0.59 | 0.176 | 0.331 |
| Separation of a loved one | 7 (8.0) | 1.20±0.79 | 5 (8.1) | 1.15±0.68 | 4 (5.0) | 1.13±0.67 | 0.684 | 0.774 |
| **Total stress load** | n.a. | 38±9 | n.a. | 41±12 | n.a. | 41±11 | n.a. | 0.078 |

P-values for differences between COVID groups were determined by performing a one-way ANOVA with Tukey-HSD post hoc test for continuous variables and a chi-square test for categorical variables.

[a]No-COVID group differs significantly from the COVID <4-week group (P<0.05). [b]No-COVID group differs significantly from the COVID >12-week group (P<0.05). [c]COVID <4-weeks group differs significantly from the COVID >12-week group (P<0.05).

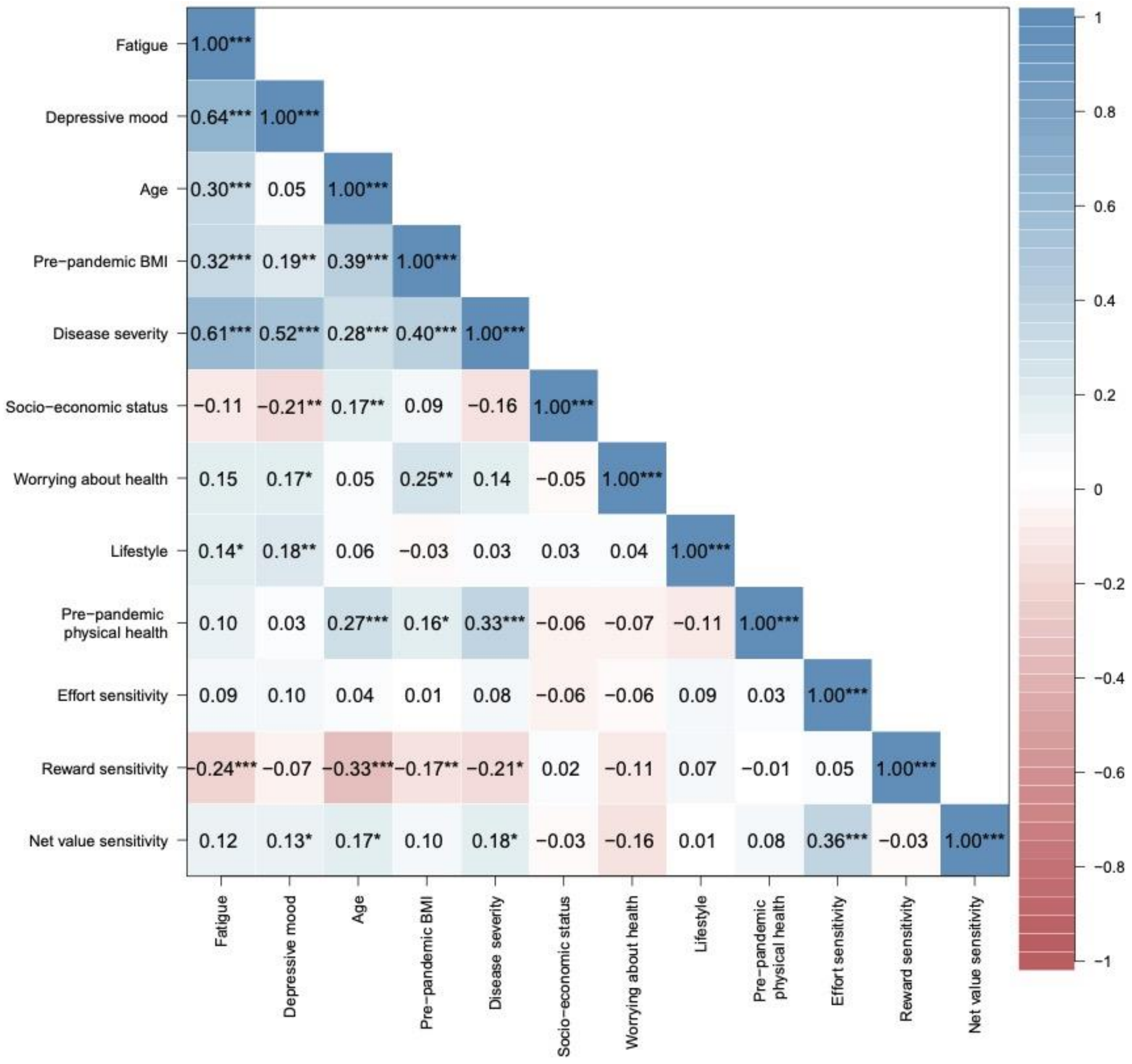


Supplementary Figure 4: Correlation matrix of predictor and outcome variables across all three groups.

*P<0.05, **P<0.01, ***P<0.001

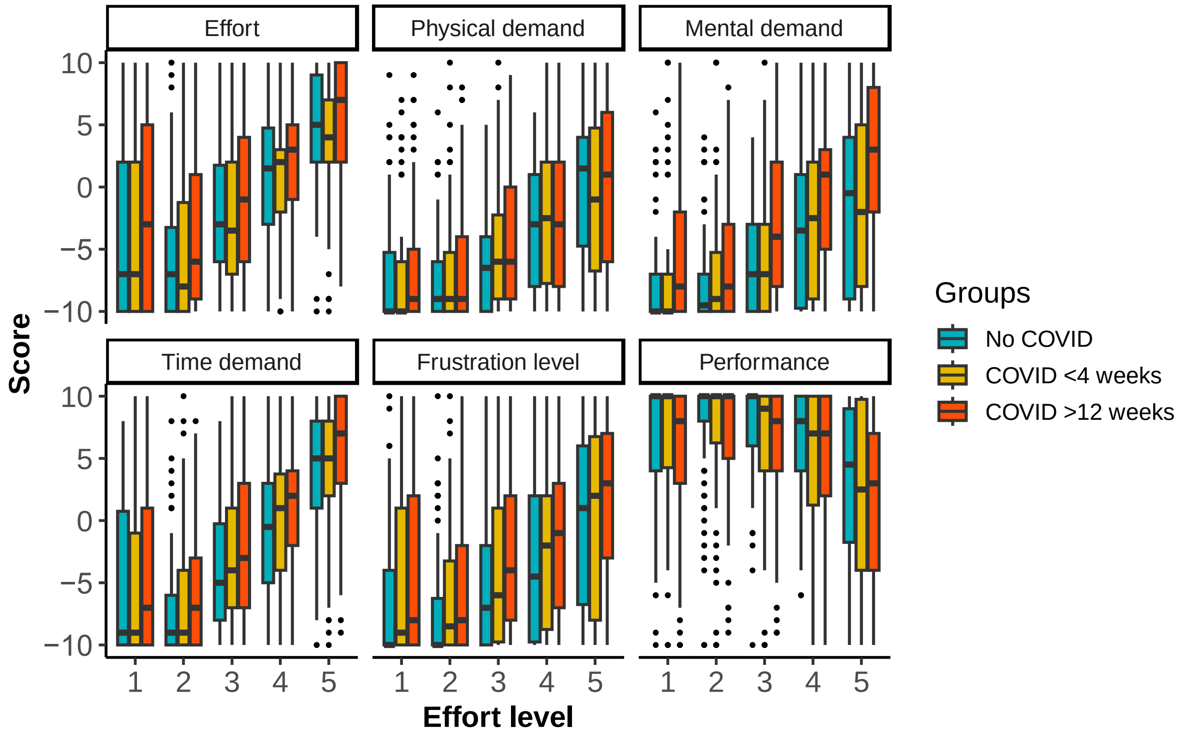


Supplementary Figure 5: NASA task load according to effort level of the effort-based decision making task.

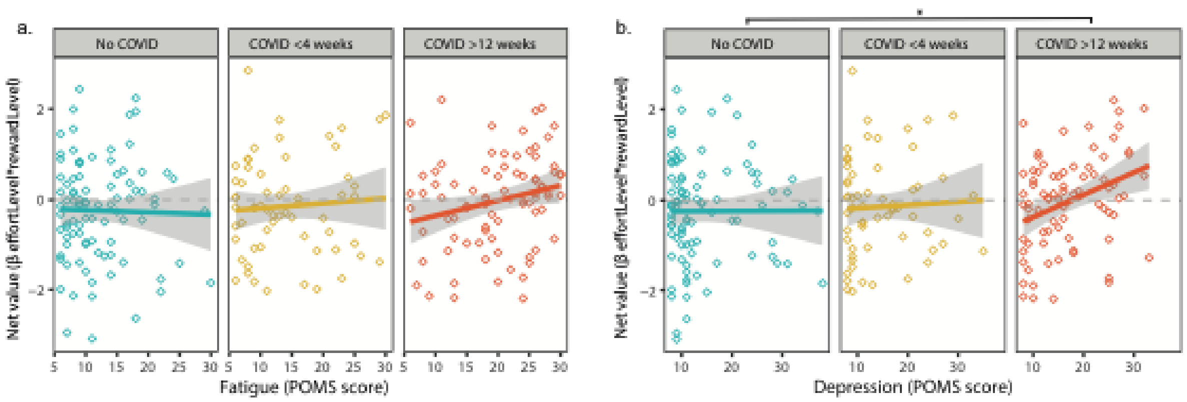


Supplementary Figure 6: The association between fatigue (a) and depressive mood (b) and the interaction between effort and reward according to group.

The regression coefficients for the interaction between effort and reward, and the differences between groups were determined and tested by mixed model binomial regression analysis.

*P<0.05

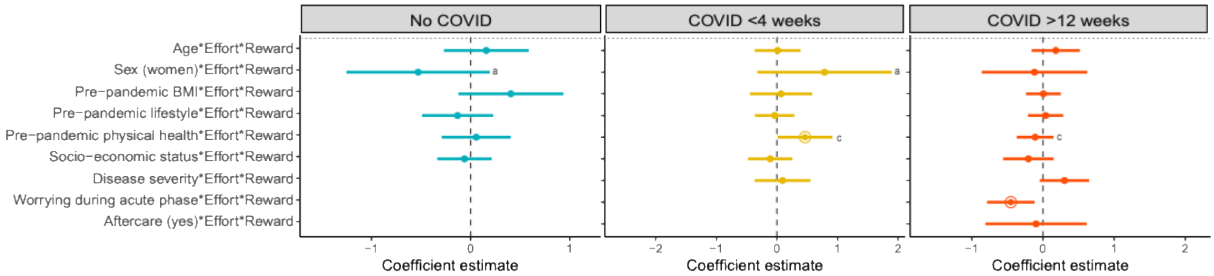


Supplementary Figure 7: Prediction regression models of the interaction between effort and reward in each group.

Regression coefficients were estimated using mixed binomial regression analysis. Continuous predictors were Z-scored. Error bars represent the 95% confidence interval. Significant regression coefficients are highlighted by a circle. A higher regression coefficient for the interaction between effort and reward represents less optimal integration of effort and reward, i.e. less choice for high effort combined with low reward options.

[a]No COVID group differs significantly from the COVID <4 weeks group (P<0.05). [b]No COVID group differs significantly from the COVID >12 weeks group (P<0.05). [c]COVID <4 weeks group differs significantly from the COVID >12 weeks group (P<0.05).

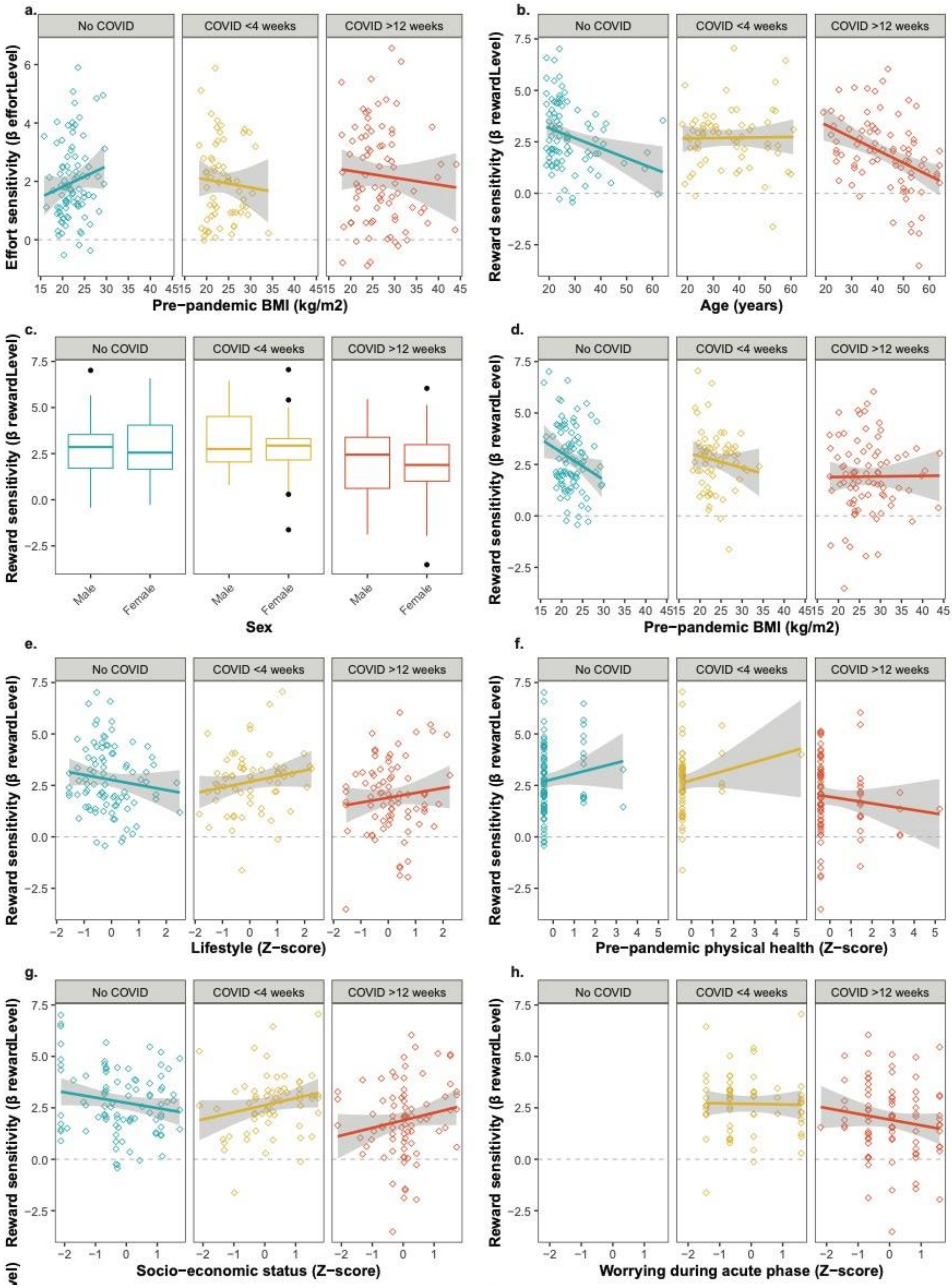
a.
b.
c.
d.
e.
f.
g.
h.
No COVID
COVID <4 weeks
COVID >12 weeks
Effort sensitivity (β effortLevel)
Reward sensitivity (β rewardLevel)
Pre−pandemic BMI (kg/m2)
Age (years)
Sex
Male
Female
Lifestyle (Z−score)
Pre−pandemic physical health (Z−score)
Socio−economic status (Z−score)
Worrying during acute phase (Z−score)

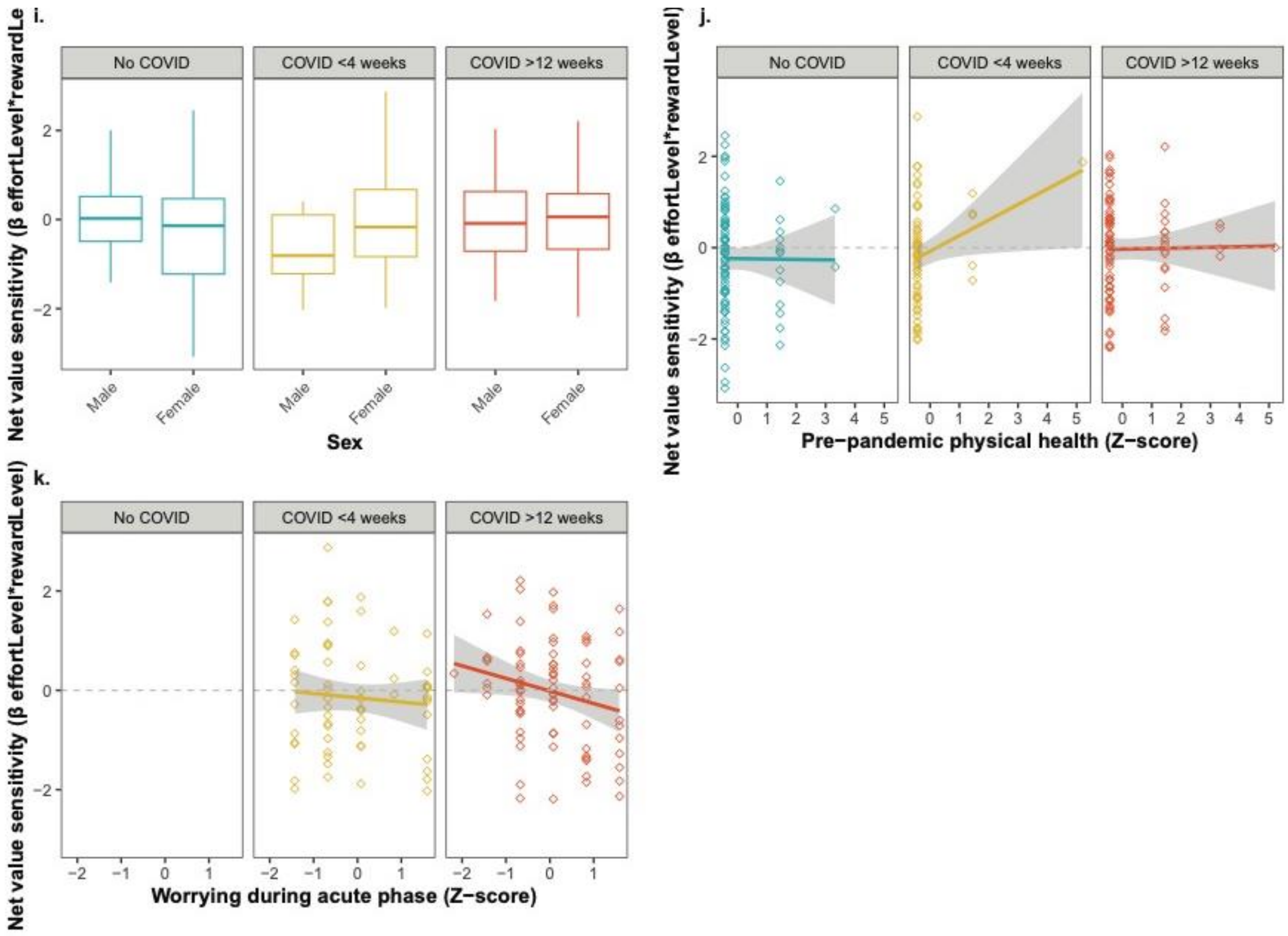


Supplementary Figure 8: Scatterplots of significant predictors and effort, reward, and net value sensitivity according to group.

Supplementary information 1:
R-codes

```
glmer(answer ~ Fatigue*Effort*Reward*Group + Sex + Age + (1 +
Effort*Reward | subject),
                 family = binomial,
                 data = data_long,
                 control=glmerControl(optimizer =
c("Nelder_Mead","bobyqa"), optCtrl=list(maxfun=1e+9)))


glmer(answer ~ Effort*Reward*group1*Sex +
                                 Effort*Reward*group*Age +
                                 Effort*Reward*group*BMI +
                                 Effort*Reward*group*Lifestyle
+

Effort*Reward*group*PhysicalHealth +

Effort*Reward*group*socioEconomicStatus +
                Effort*Reward*group*DiseaseSeverity +
                Effort*Reward*group*Aftercare +
                Effort*Reward*group*Worrying + (1 +
          Effort*Reward | subject),
                 family = binomial,
                 data = data_long,
                 control=glmerControl(optimizer =
c("Nelder_Mead","bobyqa"), optCtrl=list(maxfun=1e+9)))
```

Supplementary infomation 2:

**Text documents:**

The results of the MFI indicated that the COVID >12 weeks group experienced more general, mental, and physical fatigue as measured by the MFI in the past two weeks compared to both the COVID <4 weeks and the no COVID group. The COVID >12 weeks group also experienced more reduced motivation and reduced activity as measured by the MFI compared to the no COVID group. Results of the SF-36 indicated that the COVID >12 weeks group had a lower general health, lower physical functioning, more limitations due to physical health, and experienced more pain in the two weeks before participation compared to both the no COVID and the COVID <4 weeks group. In addition, two COVID >12 weeks group had lower social functioning in the two weeks before participation compared to the no COVID group. Emotional well-being and limitations due to emotional problems did not differ between the groups.

Figure 2 shows how often participants had physical and psychological symptoms at the time of testing compared to before the pandemic. Most common symptoms across groups were fatigue, fatigue after mild exertion, having a long recovery after exertion, concentration problems and a high heart rate. The COVID <4 weeks and COVID >12 weeks groups reported overall more physical and fatigue-related symptoms compared to participants who did not have COVID. The symptoms of feeling depressed, having a low motivation, feeling stressed, feeling anxious, rash and coughing blood did not differ between the three groups. The three groups also experienced a similar amount of pandemic-related stress (Supplementary Table 2). In addition, the COVID >12 weeks reported more symptoms during their first two weeks of their COVID infection compared to the COVID <4 weeks group (Supplementary Figure 2).
